# Phenology effects on productivity and hatching-asynchrony of American kestrels (*Falco sparverius*) across a continent

**DOI:** 10.1101/2022.01.14.476385

**Authors:** Kathleen R. Callery, Sarah E. Schulwitz, Anjolene R. Hunt, Jason M. Winiarski, Christopher J. W. McClure, Richard A. Fischer, Julie A. Heath

## Abstract

Climate-driven advances in spring can result in phenological mismatch between brood rearing and prey availability and consequently cause decreased productivity in birds. How consequences of mismatch vary across species’ ranges, and how individual behavior can mitigate mismatch effects is less studied. We quantified the relationship between phenological mismatch, productivity, and behavioral adaptations of American kestrels (*Falco sparverius*) across their breeding range in the United States and southern Canada. We obtained phenology and productivity data using nest observations from long term nest box monitoring, remote trail cameras, and community-scientist based programs. We collected data on parental incubation behavior and hatch asynchrony using trail cameras in nest boxes. Kestrels that laid eggs after the start of spring had higher rates of nest failure and fewer nestlings than earlier nesters, and effects of mismatch on productivity were most severe in the Northeast. In contrast, kestrels in the Southwest experienced a more gradual decline in productivity with seasonal mismatch. We attribute the effect of location to the growing season and temporal nesting windows (duration of nesting season). Specifically, resource availability in the Northeast is narrow and highly peaked during the breeding season, potentially resulting in shorter nesting windows. Conversely, resource curves may be more prolonged and dampened in the Southwest, and growing seasons are becoming longer with climate change, potentially resulting in longer nesting windows. We found that the onset of male incubation was negatively associated with lay date. Males from breeding pairs that laid eggs after the start of spring began incubation sooner than males from breeding pairs that laid before the start of spring. Early-onset male incubation was positively associated with hatching asynchrony, creating increased age variation in developing young. In sum, we demonstrate that American kestrels are vulnerable to phenological mismatch, and that this vulnerability varies across space. Northeastern populations could be more vulnerable to mismatch consequences, which may be one factor contributing to declines of kestrels in this region. Also, we demonstrate early onset of incubation as a potential adaptive behavior to advance average hatch date and spread out offspring demands, but it is unknown how impactful this will be in mitigating the fitness consequences of phenology mismatch.

**Highlights:** Climate-driven phenological mismatch is a growing conservation concern.

We studied phenology and productivity of an avian predator across North America.

American kestrels nesting after the start of spring had lower productivity.

Later nesting kestrels altered incubation to create hatching asynchrony.

Mismatch effects were strongest in the Northeast and may contribute to population declines.

## Introduction

Climate change is impacting the onset of spring and the duration of the growing season across temperate regions (Schwartz et al., 2006; Christiansen et al., 2011). Phenology (i.e., the timing of seasonal life events) has shifted unequally among different taxa and trophic levels, even between species that are ecologically linked (Walther et al., 2002), resulting in phenological mismatches between animal breeding seasons and food availability (Visser et al., 1998; Buse et al., 1999). In birds, mismatches between timing of offspring rearing and peak food availability may result in lower food provisioning, slower growth, poor body condition, and high nestling mortality (Buse et al., 1999; Sanz et al., 2003; Visser et al., 2006; Reed et al., 2013; Wann et al., 2019). Hence, individuals that track resource phenology more closely tend to have higher nest success and productivity than individuals that are mismatched (Forchhammer et al., 1998; Both et al., 2004; Visser et al., 2012). As climate change continues, fitness consequences of phenological mismatch may contribute to population declines, especially when the rate of spring advancement outpaces species’ capacities for adaptation (Visser et al., 2012). Therefore, it is important to understand population characteristics associated with vulnerability to phenological mismatch.

Although the evidence for consequences of phenological mismatch on productivity is widespread in birds, the effects are not homogenous across regions, even within a species. Growing seasons and climatic conditions vary widely across North America, with some regions experiencing narrow, peaked resource availability during the breeding season, and others experiencing a prolonged, dampened resource curve. The former (i.e., a short nesting window) may result in very high productivity for individuals that optimally time breeding, but may have extreme consequences for individuals that mistime breeding. Conversely, the latter (i.e., long nesting window) may result in lower productivity peaks, but mistiming effects may be less severe (Garcia-Heras et al., 2016). Indeed, seasonal declines in productivity (Garcia-Heras et al., 2016) and population declines (Both et al., 2010) are both more pronounced for species nesting in regions with strong seasonality and shorter nesting windows than those nesting in regions with weaker seasonality and longer nesting windows. Climate change effects also vary regionally, with the steepest spring warming and increases in frost free days occurring in northern and western regions (Easterling, 2002; Peterson et al., 2013), extreme temperatures and drought occurring in southern and western regions (Peterson et al., 2013), and increasing extreme precipitation events occurring in eastern regions (Kunkel et al., 2013; Huang et al., 2017). This regional variation in climate change impacts will likely cause variation in the width and flexibility of nesting windows, and potentially change resulting fitness consequences for birds.

When mismatch does occur, behavioral plasticity of individuals offers one potential mechanism for species to respond and adapt to resource limited conditions associated with mismatch. For example, initiating continuous incubation before clutch completion results in earlier average hatch date (i.e., less mismatched, Both & Visser, 2005), and staggers egg hatching dates and nestling development in a phenomenon called “hatch asynchrony” (Clark & Wilson, 1981). Producing offspring that reach their peak growth rate at different times lessens the per diem energy burden on parents during brood-rearing (Wiebe & Bortolotti, 1994; Mainwaring et al., 2014), which could be adaptive if brood-rearing is occurring under mismatched, resource-limited conditions.

American kestrels (*Falco sparverius*) are a widespread, generalist predator that breeds across much of North America. They have broad diets and prey on insects, small mammals, birds, and lizards (Smallwood & Bird, 2020), and start of spring is a good proxy for abundance of kestrel prey species (Lafage et al., 2014, Smith et al., 2017). Kestrel lay dates are advancing with earlier springs in western populations (Smith et al., 2017), but remain unchanged in eastern populations, despite advancing springs (Del Corso, 2016). The causes of geographic variation in phenological shifts, and potential fitness consequences are unknown. Further, geographic variation in population trends, specifically steeper declines in eastern populations (Smallwood et al., 2009; McClure et al., 2017), emphasize the importance of understanding population characteristics associated with vulnerability to mismatch consequences.

Our objectives were to study the consequences of American kestrel phenological mismatch on productivity across a large scale, and determine whether kestrels employ behaviors to mitigate mismatch effects. Specifically, we quantified mismatch of breeding time (i.e., lay date of first egg) relative to the start of spring, and kestrel productivity (i.e., number of fledglings produced per nest) using data from long-term monitoring projects and community science programs across North America spanning nearly 25 years. We also quantified onset of incubation and degree of hatching asynchrony using image data from remote trail cameras in nest boxes located on Department of Defense sites across the United States in 2018 – 2019. We predicted that productivity would decline as phenological mismatch increased, but that this pattern would vary spatially due to regional differences in seasonality. Finally, we predicted that laying after the start of spring would result in earlier onset of incubation behavior and lead to hatching asynchrony.

## Methods

### Clutch initiation date, nest outcome, and productivity

We obtained nest records from the Cornell Lab of Ornithology’s NestWatch program (1997 - 2018; https://nestwatch.org/) and The Peregrine Fund’s American Kestrel Partnership (AKP, 2007 – 2019; https://kestrel.peregrinefund.org/). Additionally, we collected data as part of our long-term *Southwestern Idaho Kestrel Study* (2008-2018), and the *Full Cycle Phenology Project* (2018-2019) where we monitored nest boxes on Department of Defense (DoD) installations in Washington, New Mexico, California, New York, North Carolina, and Kansas (Figure 1).

**Figure 1.**
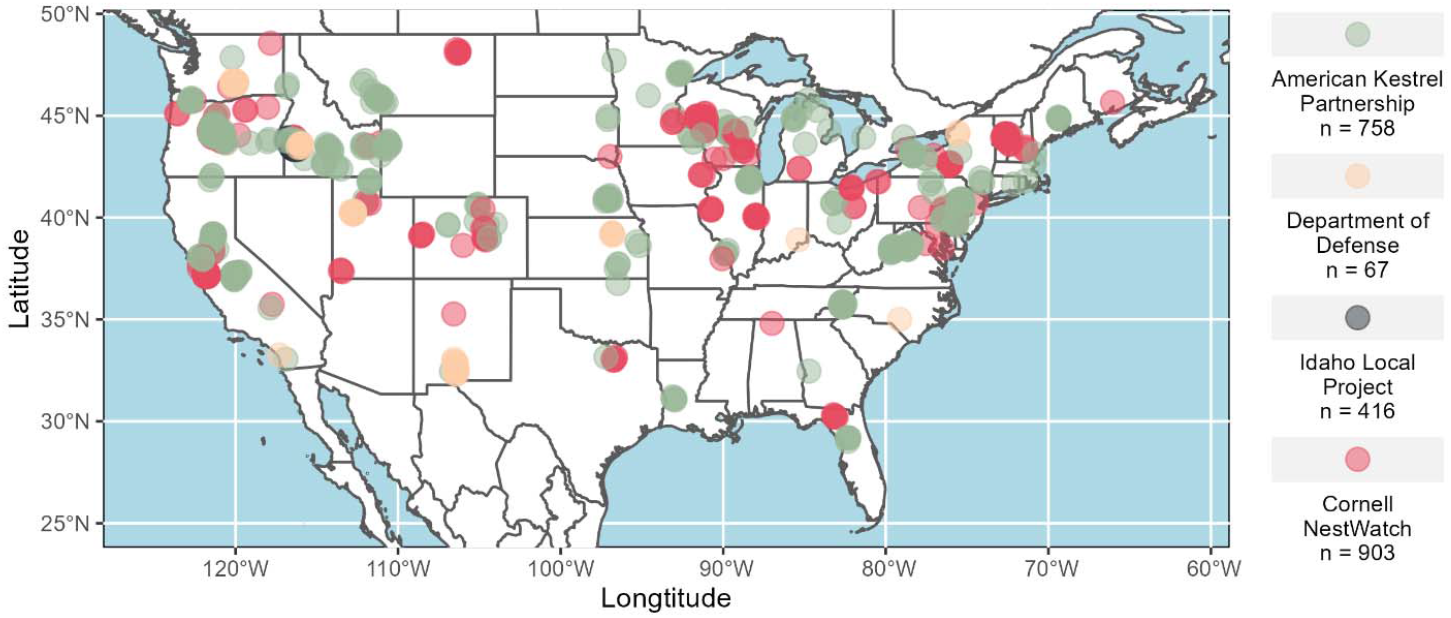
Map of American kestrel nests included in the productivity analysis (n = 2144). Each point represents one nest, and the color of the point indicates the source of data: the American Kestrel Partnership (2007-2019) (n = 758), the Full Cycle Phenology Project on Department of Defense land (2018-2019) (n = 67), the long-term monitoring site in southwestern Idaho (2008-2018) (n = 416), or Cornell NestWatch (1997-2018) (n = 903).

Data collection protocols varied depending on the monitoring program, but nest records were typically collected through repeat monitoring by community science volunteers, professional biologists, time-lapse camera imagery, or a combination of these approaches. We restricted our analysis to nests in which 1) eggs or nestlings were observed and the date on which the first egg of the clutch was laid (clutch initiation date; CI-date) was directly observed or could be reliably back-calculated from the information provided, 2) nest outcome could be assigned from nestling age or participant comments, and 3) the number of young fledged was provided. Additional details on data collection for each of these nest monitoring programs, data processing steps, and determination of breeding parameters are provided in Supplementary Material.

#### Incubation onset and hatch asynchrony

We quantified incubation onset and hatch asynchrony at a subset of DoD nest boxes equipped with cellular or non-cellular trail cameras (Spypoint, see Supplement Material) that captured complete time-lapse imagery from clutch initiation date through the end of the incubation period. Cameras were initially programmed to capture three images per day, but cellular cameras were switched remotely via the Spypoint website to take hourly images once the first egg was laid. We defined the relative onset of incubation behavior as the difference in days between clutch initiation date and the first day in which each parent began incubation (i.e., laying prone over the eggs and the majority of the eggs are covered). For hatch asynchrony, we calculated the variation in plumage-determined ages (Griggs & Steenhof, 1993) among nestlings approximately 23 – 25 days after the first egg hatched.

#### Start of spring estimates

We used extended spring-index (SI-x) models developed to predict the first-bloom dates of lilac (*Syringa chinensis* and *S. vulgaris*), and honeysuckle cultivars (*Lonicera tatarica* and *L. korolkowii*) (Schwartz et al., 2006; Rosemartin et al., 2015) to estimate the start of spring across North America. Lilac and honeysuckle first-bloom dates have been used to indicate the onset of spring, and the ubiquitous distribution of these ornamental plants allows for the meaningful comparison of spring phenology across space, time, and different biomes (Schwartz & Hanes, 2010). Further, Callery et al. (*in review*) found that American kestrel nest initiation is positively associated with SI-x. We extracted SI-x dates derived from Daymet climate datasets (Thornton et al., 2018) at the latitude and longitude of each occupied nest box per year using Google Earth Engine code modified from Izquierdo-Verdiguier et al. (2018). We created an index of phenology mismatch by calculating the difference in days between the clutch initiation date and the SI-x date.

#### Statistical analysis

We used a zero-inflated generalized linear mixed-effect model with a Generalized Poisson distribution and log-link to evaluate candidate model sets for predicting productivity in the “glmmTMB” package (Brooks et al., 2017) for R (R Core Team, 2020). This model included two sub-models: 1) zero-inflation to model probability of nest failure, and 2) conditional generalized Poisson to model count data (i.e., number of young fledged from successful nests). Each model in this candidate set included the random effect of the year. Covariates included in the conditional and zero-inflation model candidate sets for productivity were phenological mismatch, latitude, and longitude. All covariates were scaled and centered. We evaluated candidate models for the zero-inflation model with an intercept-only conditional model first. Then, we used the best supported zero-inflated model to evaluate candidate models for the conditional model.

We created gamma-distributed generalized linear models with log links to examine the relationship between within-brood variation in nestling age and the timing of the onset of incubation behavior separately for each parent. Then, we used generalized linear models with negative binomial distributions and a log link to determine if parental incubation behavior (number of days between clutch initiation date and the first day of incubation) was predicted by phenological mismatch or location. For these models we used data from both successful and unsuccessful nest attempts with complete photographic records of incubation behavior.

We compared candidate models using Akaike’s information criterion corrected for small sample size (AICc), and considered the models within 2ΔAICc to be informative (Burnham & Anderson, 2002). We estimated 85% confidence intervals for parameters in the top model to be compatible with model selection criteria (Arnold, 2010). We conducted all analyses in R (R Core Team, 2020).

### Results

We collected data from 2144 American kestrel nesting attempts that occurred between 1997 and 2019 in the contiguous US and southern Canada (Figure 1). Most kestrel nests were successful (n = 1642, 77%) and raised 1 - 7 young (mean = 3.9, standard deviation = 1.1). Egg laying dates ranged from 1 March - 14 June. On average, kestrels nested 8 days (± 0.5 days) before the start of spring.

The best zero-inflation model for predicting reproductive outcome included an interaction between phenological mismatch, latitude, and longitude (Table 1A). Kestrels were more likely to fail if they nested after the start of spring, and this effect was strongest in the northeast β_async*lat_ = 0.18, 85% CI: 0.09 – 0.27; β_async*long_ = 0.40, 85% CI: 0.25 – 0.54; β_lat*long_ = −0.33, 85% CI: −0.44 – −0.22). The best conditional model for American kestrel productivity was the additive effect of phenological mismatch β_asynct_ = −0.07, 85% CI: −0.08 – −0.05) with an interaction between latitude and longitude (β_lat*long_ = 0.0.3, 85% CI: 0.02 – 0.04; Table 1B). These results suggest that productivity was lower for successful pairs that nested after the start of spring, regardless of location. When nesting earlier relative to the start of spring, kestrels in the northeast had more young per brood than kestrels in the West and Southwest (Figure 2). However, northeastern kestrels experienced a sharper decline in productivity with increasing mismatch than kestrels from other regions included in our study (Figure 2).

**Figure 2.**
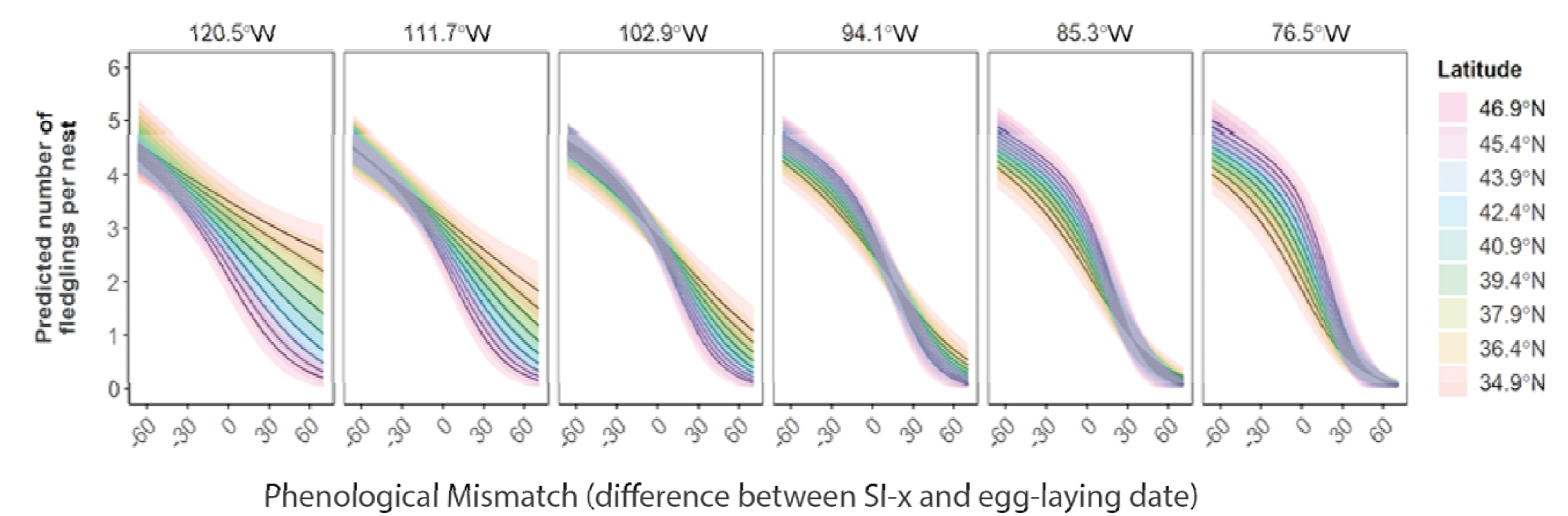
The number of fledglings produced per nest attempt was best predicted by the additive effect of phenological mismatch (the difference in days between the clutch initiation date and the extended spring index date) and the interactive effect of latitude and longitude. The number of young fledged per nesting attempt decreased as pairs laid eggs after the start of spring. This effect was strongest in the northeast, where productivity was highest then steeply declined. The lines represent the model prediction, the shaded regions are the 85% confidence intervals of the prediction, the facets display predictions at different longitudes, and the colors indicate predictions at different latitudes.

**Table 1.** Candidate models for predicting the number of American kestrel young fledged per brood. Zero-inflated generalized Poisson mixed-effect linear models included the covariates of phenological mismatch (the difference between lay date and the start of spring), longitude in °W, and latitude in °N. Zero-inflated models (A) had an intercept only conditional model. Conditional models (B) included the best model for zero-inflation (all two-interactions among mismatch, latitude and longitude). Each conditional and zero-inflation model included a random effect of year. Tables show models with weights > 0.01 and an intercept-only model, number of parameters estimated (K), ΔAICc, and model weights (*w*_i_).

| Zero-Inflation Model Formulas | K | $\Delta\text{AICc}$ | $w_i$ |
| --- | --- | --- | --- |
| Mismatch* Latitude + Latitude*Longitude + Mismatch*Longitude | 11 | 0.0<sup>a</sup> | 0.70 |
| Mismatch * Latitude * Longitude | 12 | 1.9 | 0.27 |
| Mismatch* Longitude + Latitude* Longitude | 10 | 6.4 | 0.03 |
| Intercept-only | 5 | 159.1 | 0 |

<sup>a</sup>AICc = 7040.9

B.
| Conditional Model Formulas | K | $\Delta\text{AICc}$ | $w_i$ |
| --- | --- | --- | --- |
| Mismatch + Latitude * Longitude | 15 | 0.0<sup>a</sup> | 0.64 |
| Mismatch * Latitude + Latitude*Longitude + Mismatch*Longitude | 17 | 1.9 | 0.25 |
| Mismatch * Latitude * Longitude | 18 | 3.4 | 0.12 |
| Intercept-only | 11 | 64.0 | 0 |

<sup>a</sup>AICc = 6976.9

There were 27 nests with complete photographic records of incubation to use for our analysis of incubation behavior. Male kestrels started to incubate 1 – 20 days (mean = 8.0, standard deviation = 4.2) after clutch initiation and females started to incubate 0 – 8 days (mean = 2.0, standard deviation = 2.3) after clutch initiation. Of the 27 nests, we had 16 successful nests where we measured variance in nestling age within broods. Within-brood nestling age variance ranged from 0 – 3 days (mean = 1.8, standard deviation = 1.3), was best explained by the onset of male incubation behavior, (β = −0.33, 85% CI: −0.47 – −0.17, Table 2, Figure 3), where early-onset of male incubation resulted in more asynchronous hatching, producing greater variance in nestling ages.

**Figure 3.**
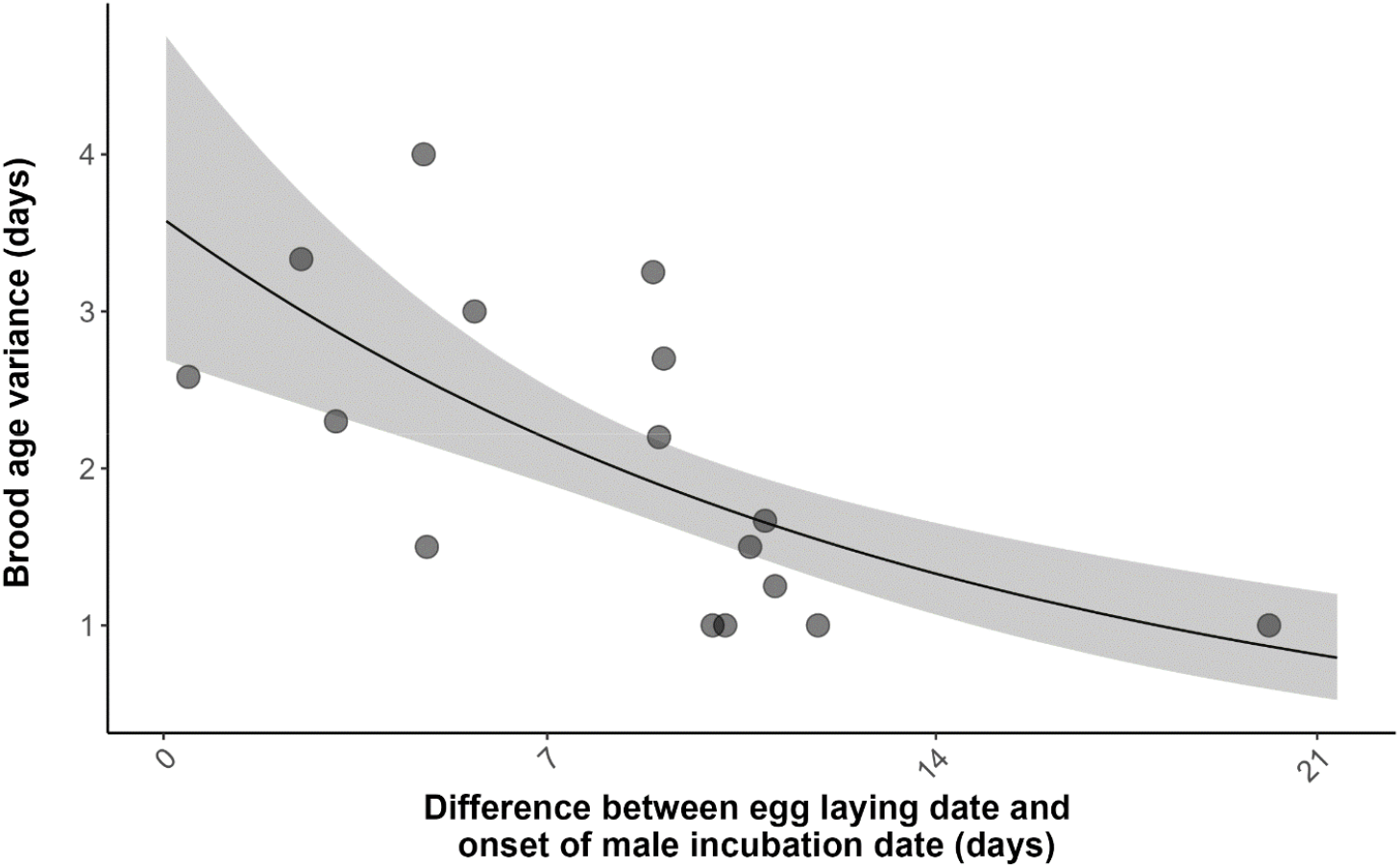
Brood age variance at fledging age was best predicted by the difference in days between the onset of male incubation and egg laying date. Each point represents a nest with complete incubation data that had at least two fledglings during the breeding seasons of 2018 (n = 8) and 2019 (n = 8). The line represents the model prediction, and the shaded region is the 85% confidence interval for that prediction.

**Table 2.** Parameter estimates, standard error, and 85% confidence intervals (LCI = lower confidence interval; UCI = upper confidence interval) from the top zero-inflation model (A), and the top conditional model (B) explaining American kestrel productivity 1997-2019 across North America. The zero-inflation models represent the probability of nest failure, whereas the conditional models predict the number of young that fledge from successful nests.

**A.**
| Parameters | Estimate | 85% LCI | 85% UCI | Std. Error |
| --- | --- | --- | --- | --- |
| (Intercept) | -1.33 | -1.58 | -1.07 | 0.18 |
| Mismatch | 0.98 | 0.85 | 1.12 | 0.10 |
| Latitude | -0.01 | -0.11 | 0.09 | 0.07 |
| Longitude | -0.29 | -0.38 | -0.19 | 0.07 |
| Mismatch * Latitude | 0.18 | 0.09 | 0.27 | 0.07 |
| Mismatch * Longitude | 0.40 | 0.26 | 0.54 | 0.10 |
| Latitude * Longitude | -0.33 | -0.45 | -0.22 | 0.08 |

**B.**
| Parameters | Estimate | 85% LCI | 85% UCI | Std. Error |
| --- | --- | --- | --- | --- |
| (Intercept) | 1.365 | 1.35 | 1.38 | 0.01 |
| Mismatch | -0.067 | -0.08 | -0.05 | 0.01 |
| Latitude | 0.001 | -0.01 | 0.01 | 0.01 |
| Longitude | 0.004 | -0.01 | 0.01 | 0.01 |
| Latitude * Longitude | 0.034 | 0.02 | 0.04 | 0.01 |

The onset of male incubation was best predicted by the additive effects of phenological mismatch (β = −0.33, 85% CI: −0.51 – −0.14) and latitude (β = −0.34, 85% CI: −0.52 – −0.16, Table 3, Figure 4). Males from breeding pairs that laid eggs after the start of spring began incubating the eggs sooner after clutch initiation than males from breeding pairs that laid before the start of spring. Males breeding in southern regions were the most likely to delay start of incubation, whereas males breeding in northern regions were most likely to initiate incubation soon after the first egg was laid.

**Figure 4.**
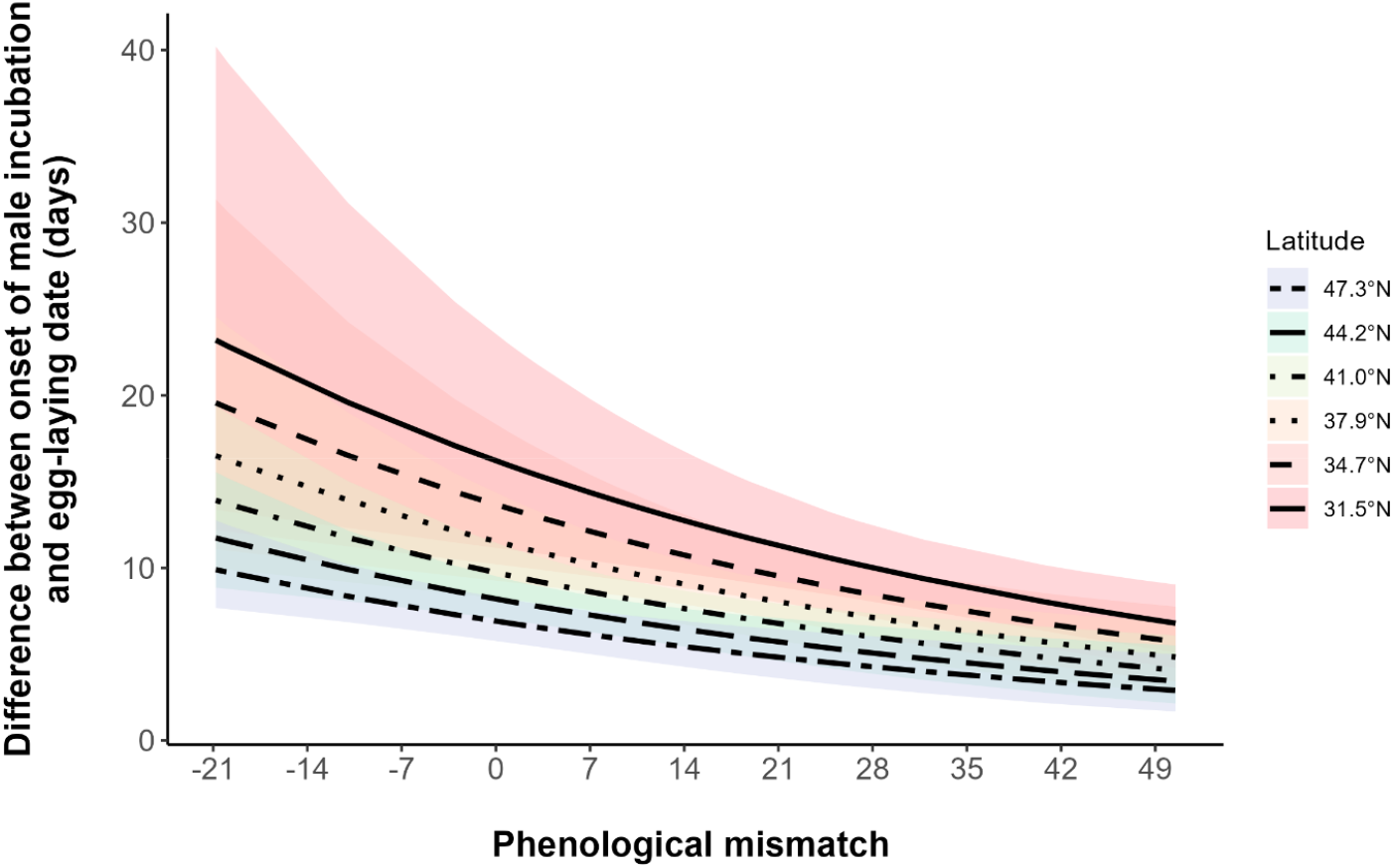
The predicted relationships between the phenological mismatch and the onset of male incubation relative to lay date (days) for different latitudes. As mismatch and latitude increase, the difference between lay date and the onset of male incubation behavior decreases. The earlier onset of male incubation behavior is a predictor for increased age variance of the nestlings, and an indicator of hatching asynchrony. The line represents the model predictions, the shaded regions are the 85% confidence interval for each prediction, and the line type of each prediction and the color surrounding it represent predictions at different latitudes.

**Table 3.** Candidate set of models to explain age variation within broods of American kestrels. Covariates included the difference between the incubation onset and laydate for male and female American kestrels. Tables show models, number of parameters estimated (K), ΔAICc, and model weights (*w*_i_).

| Candidate model | K | $\Delta\text{AICc}$ | $w_i$ |
| --- | --- | --- | --- |
| Male Incubation | 3 | 0.0<sup>a</sup> | 0.79 |
| Male Incubation + Female Incubation | 4 | 3.2 | 0.16 |
| Intercept-only | 2 | 6.2 | 0.04 |
| Female Incubation | 3 | 8.6 | 0.01 |

<sup>a</sup>AICc = 40.3

**Table 4.** Candidate set of models to explain the onset of male incubation in American kestrels. The covariates included are phenological mismatch, latitude, and longitude. The table shows model, number of parameters estimated (K), ΔAICc, and AICc weights (*w*_i_).

| Candidate model | K | $\Delta\text{AICc}$ | $w_i$ |
| --- | --- | --- | --- |
| Mismatch + Latitude | 4 | 0.0<sup>a</sup> | 0.42 |
| Mismatch + Latitude + Longitude | 5 | 0.7 | 0.29 |
| Intercept-only | 2 | 3.2 | 0.09 |

<sup>a</sup>AICc = 150.5

## Discussion

We show the negative consequences of phenological mismatch on productivity of American kestrels, regional variation in strength of effects, and potential behavioral adaptations to mismatch. The negative effects of mismatch were strongest in the Northeast, where kestrels experience highly peaked resource availability. The effects of phenological mismatch may be mitigated by males advancing onset of incubation of later laid nests, which resulted in hatching asynchrony and variation in nestling ages.

Consistent with previous literature (Goodenough et al., 2010; Bowers et al., 2016; Taylor et al., 2021), we showed that mismatch, specifically laying eggs after the start of spring, decreased both nest success and productivity of American kestrels across their range. Geographic variation in strength of mismatch effects (i.e., stronger effects in the Northeast) may be related to variation in growing seasons and climate change impacts (Both et al., 2010; Garcia-Heras et al., 2016). Growing seasons in the Northeast have a higher but narrower peak in productivity in the spring than in the West, where green-up is less peaked and more heterogeneous (Callery et al., *in review*). This may explain why “on-time” nesters in the Northeast have higher productivity peaks, but face steeper productivity declines due to mistimed breeding (i.e., shorter nesting window), compared to kestrels in regions where less-peaked, but prolonged growing seasons allow more flexibility in breeding time (i.e., longer nesting window). Similarly, Garcia-Heras et al. (2016) found that black harriers (*Circus maurus*) in regions with short breeding seasons had more pronounced seasonal declines in productivity than those in regions with long breeding seasons. The nesting window for American kestrels in the Northeast is also constrained by the increasing frequency of extreme precipitation events in winter and early spring (Huang et al., 2017) which can delay migrant arrival time (Powers et al., 2021), and cause decreased foraging ability, prey availability, and lower productivity in early arriving kestrels (Olsen & Olsen, 1992; Dawson & Bortolotti, 2000; McDonald et al., 2004). These climatic conditions are creating an increasingly inflexible and narrow time window (e.g. ~2 month lay date range in New Jersey; Del Corso, 2016) within which northeastern kestrels can breed without experiencing a decrease in productivity. Conversely, in western North America winters are becoming milder, which has been associated with shorter kestrel migration distances (Heath et al., 2012), northward shifts in wintering distributions (Paprocki et al., 2014), and earlier breeding (Heath et al., 2012). The onset of spring is also advancing more rapidly in the Mountain West than anywhere else in our study region (Schwartz et al., 2006; Allstadt et al., 2015), and farmers are advancing the start of their planting season (Christiansen et al., 2011; Smith et al., 2017), resulting in wider prey peaks and long nesting windows in the West (e.g. ~ 4 month lay date range in southwestern Idaho; Steenhof & Peterson, 2009). Less is known about kestrel demographics in the Southwest, but southern populations are less migratory (Smallwood & Bird, 2020), and experience mild winters and long nesting windows (e.g. ~ 4 month lay date range in northwest Texas: Mullican, 2018), which may make them less vulnerable to mismatch. Lastly, differing adaptive capacity of distinct genetic groups (i.e., East, West, Alaska, Texas, and Florida) and migratory phenotypes (i.e., resident or migratory) may contribute to regional discrepancies in mismatch effects (Smallwood et al., 2009; Heath et al., 2012; Ruegg and Brinkmeyer et al., 2021).

We documented early onset incubation, and resulting hatch asynchrony, as a potential adaptation to phenological mismatch in American kestrels. Similar to other species, we demonstrated onset of continuous incubation as a mechanism for producing hatching asynchrony in American kestrels (Clark & Wilson, 1981). Onset of male incubation was associated with phenological mismatch of clutch initiation with the start of spring and latitude. Males from breeding pairs that laid eggs late relative to the start of spring began incubating shortly after the first eggs were laid, which advanced the average hatch date of later broods and increased nestling age variance, consistent with the “hurry-up” hypothesis (Clark & Wilson, 1981). This relationship between mismatch and incubation onset suggests that hatching asynchrony may be an adaptation to low food resources resulting from sub-optimal breeding timing. Indeed, asynchronous hatching in American kestrels has been documented more frequently in years of food scarcity, and asynchronous kestrel broods need less provisioning per day than synchronous broods (Wiebe & Bortolotti, 1994). Our results also showed that males breeding at higher latitudes were more likely to initiate incubation earlier in the laying order than those at lower latitudes. Selection pressure for optimal breeding time is stronger at higher latitudes compared to lower latitudes (Shave et al., 2019) due to narrow, peaked resource availability. However, because of constraints of long-distance migration (Heath et al., 2012), incomplete knowledge of breeding ground conditions (Samplonius et al., 2018), and early season weather stochasticity, high latitude breeders may be unable to adjust their breeding time to match resource peaks (Hurlbert & Liang, 2012; Powers et al., 2021). Indeed, American kestrels breeding at high latitudes (long-distance migrants) did not adjust arrival time according to spring temperatures, whereas low latitude breeders (short-distance migrants) arrived earlier in earlier springs (Powers et al., 2021). Hence, early onset incubation to induce hatch asynchrony could be an alternative strategy for high latitude kestrels to cope with mismatch.

Interestingly, we found that male incubation, but not female incubation, predicted onset of continuous incubation in American kestrels. Although females tended to start incubating early in the egg laying sequence, the contribution of the male may have provided the additional incubation necessary to stimulate embryo development (Nilsson, 1993). Alternatively, our methods may have more accurately measured male incubation behavior because it was unlikely males would lay on the eggs for any other purpose but incubation, whereas a female laying eggs could be confused for a female in incubation posture.

In summary, nesting after start of spring has negative consequences for productivity of American kestrels across their range. With climate change increasing the amount of mismatch between trophic levels, the demonstrated fitness consequences of mismatch could result in population-level effects for American kestrels. The steepest mismatch effects (i.e., declines of nest success and productivity) were seen in the Northeast, which is particularly concerning in light of declining trends in northeastern populations, and constraints on phenology shifts (i.e., short growing season and early inclement weather). Although early onset of incubation may be an adaptive behavior to advance average hatch date and spread out offspring demands, it is unknown how impactful this will be in mitigating the fitness consequences of phenology mismatch.

## Acknowledgements

Funding for this project was provided by a grant from the Strategic Environmental Research and Development Program (SERDP) of the US Department of Defense (US DoD; Award Number: RC2702) and Scott Radford and other generous donors to the Boise State and Peregrine Fund Adopt-A-Box Program. We are grateful for the community scientist partners who contributed data. We thank principal scientist Robyn Bailey and Cornell NestWatch for contributing data to this analysis. We thank our wonderful field team: K. Callegari, K. Dreher, E. Hirsh, A. Johns, H. McCaslin, C. Pozzanghera, S. Ranck, C. Rankin, A. Santiago, and S. Shively, and partners T. Booms, P. Jeurgens. We thank our partners on DoD installations: W. Berry, J. Bolsinger, P. Cutler, A. Dankert, R. Felix, J. Ferrer-Perez, T. Filkins, M. Gagnier, J. Haddix, M. Houck, K. Hyde, K. Karssen, R. Knight, C. Leingang, J. Mangelinckx, D. Moon, K. Murbock, M. Parks, S. Phillips, C. Plimpton, D. Rees, J. Robb, B. Rossi, J. Schillaci, A. Schultz, S. Stratton, S. Sullivan, and K. White.

