## Supplementary Information for "Phenology effects on productivity and hatching-asynchrony of American kestrels (*Falco sparverius*) across a continent"

Supplementary Material

**Methods for monitoring nest contents**

*Cornell’s NestWatch*

NestWatch (<https://nestwatch.org/>) is a long-term community science project administered by the Cornell Lab of Ornithology that engages participants in monitoring the breeding biology of North American birds (Phillips & Dickinson, 2009). NestWatch asks volunteers to observe nests 1–2 times per week, although the protocol is flexible (Phillips et al., 2007). Specifically, NestWatch volunteers use standardized data forms to record nest metadata (e.g., species, year, unique site names, location, and nest substrate) and breeding parameters for each nest visit. These include date and time, numbers of eggs, live young, and dead young for host and brood parasite species, and descriptions of adult activity, nest maintenance activity, and developmental stages of nests and young via standardized codes. In addition, volunteers are encouraged to provide a summary of the host species’ estimated first egg, hatch and fledge dates, maximum clutch size, numbers of unhatched eggs, live young, and fledged birds, and any anecdotal evidence for why they believe a nest was successful or failed. Volunteers enter their nest observations online after each visit or at the end of the breeding season. The NestWatch database uses built-in error checks and filters (e.g., number of young cannot exceed the maximum clutch size) to ensure accurate data entry by participants (Phillips & Dickinson, 2009).

*American Kestrel Partnership*

The Peregrine Fund’s American Kestrel Partnership (AKP; <https://kestrel.peregrinefund.org/>) coordinates a network of community and professional scientists monitoring nest boxes across the western hemisphere, with the goal of addressing long-term kestrel population declines. Initiated in 2012, the AKP enlists volunteers to install and monitor nest boxes and to contribute their data to a centralized database. AKP partners submit characteristics for each nest box they monitor (e.g., nest box ID, geographic coordinates, substrate, orientation), date and time of each visit, and corresponding nest contents (e.g., count of adults, eggs, live and dead nestlings, nest box use by other species). If possible, the oldest nestling is aged using plumage characteristics according to a photographic guide of kestrel nestling development (Klucsarits & Rusbuldt, 2007). The AKP protocol encourages partners to check nests once every other week beginning in late winter or early spring (in North America, early March). At minimum, partners are asked to make at least one more visit within 30 days to check for nestlings. If eggs are still present on this second visit, partners are then asked to return again within 30 days. This nest check interval guarantees at least one visit when there are eggs and one visit when there should be nestlings present. Partners are encouraged to enter nest data on the same day that observations are made, or after each breeding season.

*Full Cycle Phenology Project*

The Full Cycle Phenology Project (FCPP; <https://fullcyclephenology.com/>) seeks to understand the impacts of climate change on American kestrel phenology and demographics in North America. FCPP established a network of 284 nest boxes at 13 Department of Defense (DoD) installations and partner sites. At each DoD site, nest monitoring began two weeks prior to the earliest clutch initiation date records for kestrels in the region. Depending on the study site, nests were monitored remotely via cellular trail cameras (Spypoint Link-Evo, Victoriaville, Québec, CA) located inside nest boxes, monthly in-person visits by DoD biologists to nest boxes equipped with non-cellular trail cameras (Spypoint Force-10, Victoriaville, Québec, CA and Reconyx Hyperfire 2, Holmen, Wisconsin, USA), or bi-weekly in-person visits to nest boxes without trail cameras. Cameras were placed inside a false lid located at the top of the nest box, with the camera’s field-of-view aimed toward the bottom of the box. Non-cellular and cellular cameras were programmed on time lapse mode to record photos three times per day (3:00, 9:00, 15:00), and were switched to hourly intervals once the first egg was detected in images or in-person visits. During in-person visits and time-lapse imagery review, nest contents were assessed by determining the presence, sex, and incubation behavior of adults, and the number of eggs and nestlings. Time-lapse imagery was reviewed and annotated at the end of each breeding season using the ViXeN software package (Ramachandran & Kadambari, 2018). In addition to regularly-scheduled visits and time-lapse imagery, additional visits were made on or near the estimated hatch date (30 days after a complete clutch was laid) to confirm hatching, and again prior to fledging (23-25 days after first egg hatched) to band and record the age of nestlings.

*Southwestern Idaho Kestrel Study*

The breeding biology of American kestrels in southwestern Idaho (43°N 116°W) has been monitored as part of a long-term study led by researchers at the US Geological Survey and Boise State University (Steenhof & Peterson, 2009; Anderson et al., 2016; Smith et al., 2017). The study area includes approximately 90–130 nest boxes (depending on the year; Smith et al., 2017) located in rural, residential, and agricultural areas near Boise, Kuna, and Meridian, Idaho. Beginning in March, nest boxes are visited every 7–10 days to determine occupancy and nest initiation date. After capturing adults, nest boxes are checked again on estimated hatch dates, when the oldest nestling is ~10 days old, and again at ~25 days, prior to fledging. Nest contents (number of eggs, live and dead young) are recorded during each visit. Nest failure is determined based on the presence or absence of eggs, nestlings, or adult birds (Strasser & Heath, 2013).

**Data curation**

*NestWatch*

We restricted our sample of NestWatch nests to attempts with a known clutch initiation date, hatch date or fledge date, and known outcomes (either success or failure).

*American Kestrel Partnership*

A primary objective of the AKP is determining the drivers of nest box occupancy, and submitting records of unoccupied boxes is highly encouraged. As a first step, we discarded nests that did not contain kestrel eggs or nestlings. We then restricted our sample to nests within the US and Canada and removed nests occupied by other species. We also excluded nest attempts that included nestling observations only and no age reported (which prevented back-calculating clutch initiation date), and attempts with a single egg observation (which did not contain nest outcome/productivity information). We conducted an additional manual screening of the data to remove nest records containing unclear observations (e.g., the appearance of multiple attempts entered per record) or comments that indicated problematic data entries. Finally, we retained nests during this step that had some indication of success or failure (i.e., ≥ 25-day-old nestlings, anecdotal evidence of fledging or predation), and provided a count of nestlings or fledglings from the nest record or comments used to determine nest outcome.

*Full Cycle Phenology Project*

The majority of the FCPP’s nest boxes were equipped with cellular or non-cellular trail cameras, generally allowing direct observation of clutch initiation date, nest success, and productivity. We discarded nests that had low frequency or no time-lapse imagery due to camera failures, and infrequent in-person visits that prohibited reliable estimation of phenology and nest outcome.

*Southwestern Idaho Kestrel Study*

We restricted our sample of southwestern Idaho nest records to attempts with a known clutch initiation date and known outcomes (either success or failure).

**Clutch initiation date, nest outcome, and productivity determinations**

*NestWatch*

We relied on user-specified fields in the NestWatch database to determine clutch initiation dates and nest outcome (Phillips & Dickinson, 2009). To assign clutch initiation date, we used the first egg date (FIRST_LAY_DT) field. If the first egg date contained an NA value, we back-calculated from the hatch (HATCH_DT) or fledge date (FLEDGE_DT) assuming 1 egg laid every other day, 30 days for incubation, and 30 days until fledging (Bird & Palmer, 1988; Anderson et al., 2016). We used the ‘s’ or ‘f’ prefix in the OUTCOME_CODE_LIST field to assign nest success or failure, respectively. If a nest was successful, we set productivity equal to the YOUNG_HOST_TOTAL_ATLEAST field value; otherwise, productivity was set to zero.

*American Kestrel Partnership, Full Cycle Phenology Project, & Southwestern Idaho Kestrel Study*

We estimated the clutch initiation date for each nesting attempt in several ways. When we discovered an incomplete clutch of eggs in a nest box, the clutch initiation date was calculated by subtracting the number of eggs in the clutch multiplied by two from the date that the clutch was discovered (AKP, FCPP, Southwestern Idaho Kestrel Study; Anderson et al., 2016), because kestrels tend to lay 1 egg every other day (Bird & Palmer, 1988). For nest attempts discovered with a complete clutch or already hatched, we used nestling age as determined by plumage characteristics (Griggs & Steenhof, 1993; Klucsarits & Rusbuldt, 2007) to back-calculate the clutch-initiation date by subtracting the age of the most mature nestling, 30 days for incubation, and twice the number of nestlings from the hatching date. For FCPP nest boxes with functioning cameras, we directly measured clutch initiation date from time-lapse imagery. For nest attempts in which the view of the nest box floor was obstructed, we relied on in-person nest visits and used the back-calculation methods described above.

We used the following criteria to determine nest outcome: successful nests produced at least one ≥ 25-day-old nestling (85% of fledging age; Strasser & Heath, 2013) as indicated by ageing during nest visits or from time-lapse imagery (FCPP), or based on anecdotal evidence of nest success or failure provided by volunteers (AKP). Finally, the number of young fledged from successful nests was estimated using the count of live nestlings when at least one ≥ 25-day-old nestling was present or from participant comments (AKP).
